# High variability of migration strategies in a re-established Trumpeter Swan population

**DOI:** 10.1101/2024.06.06.597790

**Authors:** David W. Wolfson, Randall T. Knapik, Anna Buckardt Thomas, Tyler M. Harms, Laura J. Kearns, Brian W. Kiss, Timothy F. Poole, Drew N. Fowler, Taylor A. Finger, Sumner W. Matteson, John J. Moriarty, Tiffany Mayo, Margaret Smith, Christine M. Herwig, David E. Andersen, John R. Fieberg

## Abstract

**Background:** The Interior Population (IP) of trumpeter swans (*Cygnus buccinator*), formerly extirpated by market hunting, was re-established in eastern North America by releasing individuals from both migratory and non-migratory populations. Their current annual movement patterns are largely unknown. Our goal was to describe their seasonal movements and quantify the proportion of the IP that is migratory, the extent and phenology of seasonal movements, and associations between movement patterns and breeding status and breeding location.

**Methods:** We deployed 113 GPS-GSM transmitters on IP trumpeter swans in six U.S. states and one Canadian province across the current IP breeding range. Using data from 252 ‘swan-years’, we estimated dates of migration events by segmenting the annual cycle using piecewise regression models fit to each yearly time-series of displacement from the breeding site. We fit a latent-state model to characterize population-level associations between breeding latitude and maximum extent of migration, and linear mixed models to quantify associations between individual characteristics (e.g., breeding status, sex) and migration phenology.

**Results:** At the individual level, 59% of swans moved to distant non-breeding-period areas (long-distance migration, defined as moving >100 km from the breeding site), 16% exhibited regional migration (25-100 km from breeding site), 19% exhibited non-migratory but local movements (<25 km from breeding site), and 6% exhibited multiple migration strategies. Swans breeding at more-northern latitudes departed their territories earlier in autumn and returned later in the spring than those breeding at more southern latitudes. Breeding swans departed later in the autumn than non-breeders, but breeding status did not have a strong association with arrival in the spring.

**Conclusion:** IP trumpeter swans are partial migrants, with a continuum of strategies each year, from local movements to long-distance migration. Much of the variability in movement patterns was related to factors tied to natural history demands (e.g., breeding status) and response to environmental conditions (e.g., through associations with breeding latitude).

## Introduction

Migration is a behavioral mechanism widely used by all major vertebrate groups (e.g., birds, mammals, fishes, reptiles, amphibians) that allows individuals to track seasonal availability of resources to increase survival and, by optimizing migration phenology, maximize long-term fitness [1–4]. Despite its prevalence as an ecological process and a large body of research involving migration, how population-level migratory traditions are established is not well understood [5]. In part, that lack of understanding is due to challenges associated with making population-level inference from observations of individuals, quantifying migratory movements along a continuum of variability, and the relative scarcity of successful reintroductions of formerly rare migratory species [6,7].

Similar to other large, long-lived avian species such as geese, cranes, and storks, adult swans provide care for their young during the first year of life, providing access to food and protection, and guiding them on their first migration cycle [8,9]. As a consequence, cultural transmission during the first year is thought to be the primary mechanism that dictates the learned migratory strategy used in subsequent years [10,11]. Although this transfer of information is an effective mechanism for preserving migratory patterns through generations, how these trends become established after a population is reintroduced on a landscape from which it had previously been extirpated is unclear. Jesmer et al. [12] found that newly translocated ungulate populations initially lost their migratory tendencies, and it took many generations to re-establish such patterns, but that migratory trends eventually stabilized.

Trumpeter swans (*Cygnus buccinator*), the largest waterfowl species in North America, were widespread throughout much of the continent prior to European colonization [13]. Due to widespread hunting for meat, skins for powder puffs, and feather quills for writing instruments, trumpeter swans were nearly extirpated in the lower 48 U.S. states and southern Canada, and reached an estimated low of 70 individuals in the 1930s [14]. In 1935, low numbers of trumpeter swans led to the establishment of Red Rock Lakes Migratory Waterfowl Refuge (renamed Red Rock Lakes National Wildlife Refuge [RRLNWR] in 1961) in the convergence of Montana, Wyoming, and Idaho, which was the last vestige of a breeding swan population in the United States (outside of Alaska) [15]. As trumpeter swan numbers at RRLNWR started to rise, this population became a source for individuals introduced in other parts of their historical breeding range. Trumpeter swans from RRLNWR were translocated to several U.S. states to augment the abundance and distribution of the diminished Rocky Mountain Population (RMP) in the inter-Mountain West and to restore the Interior Population (IP), which had been extirpated in eastern North America [15]. In the 1950s, aerial surveys in central Alaska revealed another population of trumpeter swans (the Pacific Coast Population [PCP]), and this group subsequently provided the majority of swans used for reintroductions within the IP [16,17]. An important distinction between these source populations is that PCP swans breeding in Alaska migrate to British Columbia and the northwestern United States each winter, whereas RMP swans from RRLNWR are non-migratory [18,19].

Estimates of both IP abundance and breeding distribution have increased dramatically since reintroductions began in the 1960s [20]. Trumpeter swans currently breed throughout most of the western Great Lakes region, including in Manitoba, Minnesota, Iowa, Ontario, Wisconsin, Michigan, and Ohio. Beyond estimates of population size and trends, however, there is relatively little recent information about their ecology, including seasonal movements and migration patterns. For example, it is not known what proportion of the IP remains resident on their breeding range during the non-breeding period, the extent (i.e., distance) of movement for those swans that do leave their breeding territories, the timing of migration (e.g., autumn departure, spring arrival), or the magnitude of intra- and inter-individual variability in migration behavior. A more comprehensive understanding of when and where IP swans move throughout the annual cycle, including any differences related to breeding status or latitude will better elucidate habitat requirements and inform optimal study design for surveys to index abundance.

Many factors likely influence an individual swan’s decision to leave its breeding territory after the breeding period, the distance migrating individuals travel, and the degree of among-individual variability in movements for swans breeding near each other; these factors include life history requirements, knowledge transfer of migratory traditions, and inherent genetic control of migration [21–23]. Arriving early in the spring allows individuals to re-establish and defend a breeding territory and lay and incubate eggs early during the breeding period when nest success and survival of offspring is generally highest [24–26]. Similarly, there are advantages to staying on breeding territories longer in the autumn to allow cygnets time to learn to fly and develop sufficient fat reserves to migrate to areas used during the non-breeding period [27,28]. Survival rates of migrant birds are typically lowest during the non-breeding period of the annual cycle due to challenges associated with navigating relatively unfamiliar landscapes and the high energetic demand of migration [29,30]. Yet, by migrating to a more-temperate area during the non-breeding period, swans can increase their access to food and other resources that allow them to avoid harsh winter conditions in their breeding territories, thereby balancing potential costs of migration with the benefits of increased resource availability [21,31].

It is likely that the drivers of migration vary within the IP based on the location of a swan’s breeding territory. Cues for migration can include declines in food availability, which may also be affected by density-dependent intra-specific competition [32]. Swans that spend the summer at different latitudes experience varying environmental conditions that may influence arrival in spring, such as the timing of ice melt and vegetative greenness [33,34], and departure in the autumn, such as ice formation on shallow wetlands and plant senescence [35,36]. Swans breeding farther from the equator contend with shorter growing seasons and greater pressures for offspring to sufficiently develop flight before environmental conditions dictate migrating to non-breeding areas [37]. The ‘push’ to avoid harsh winter conditions varies substantially throughout the IP breeding range, with concurrent implications for decisions related to migration propensity and timing. For many wildlife species, some individuals that migrate each year whereas others are residents (i.e., partial migration), which can result in higher overall fitness [23,38,39]. We expected the propensity for trumpeter swans to migrate to be higher in more northern latitudes.

To better understand current movement ecology of the re-established IP trumpeter swans, we marked a sample of swans with GPS-GSM transmitters and monitored their movements over multiple annual cycles. Our primary goal was to better understand how a re-established population of trumpeter swans derived from different source populations uses a novel landscape. Our specific objectives were to quantify 1) the proportion of the IP that is migratory and the extent of migratory movements, 2) migration phenology (e.g., timing of autumn departure and spring arrival), and 3) associations between annual movement patterns and breeding status and breeding location.

## Methods

### Study Area

We captured IP swans from 2019–2022 throughout their current breeding distribution [20] during the breeding period, plus 4 swans captured in Arkansas during the non-breeding period (Fig. 1). We deployed transmitters on IP trumpeter swans as far north and west as southern Manitoba (51.1° N, 99.7° W), as far south as Arkansas (35.5° N, 91.9° W), and as far east as Ohio (41.4° N, 80.7° W). Capture locations occurred in a mix of Laurentian Mixed Forest, Prairie Parkland, Eastern Broadleaf Forest, and Aspen Parklands [40].

**Figure 1:**
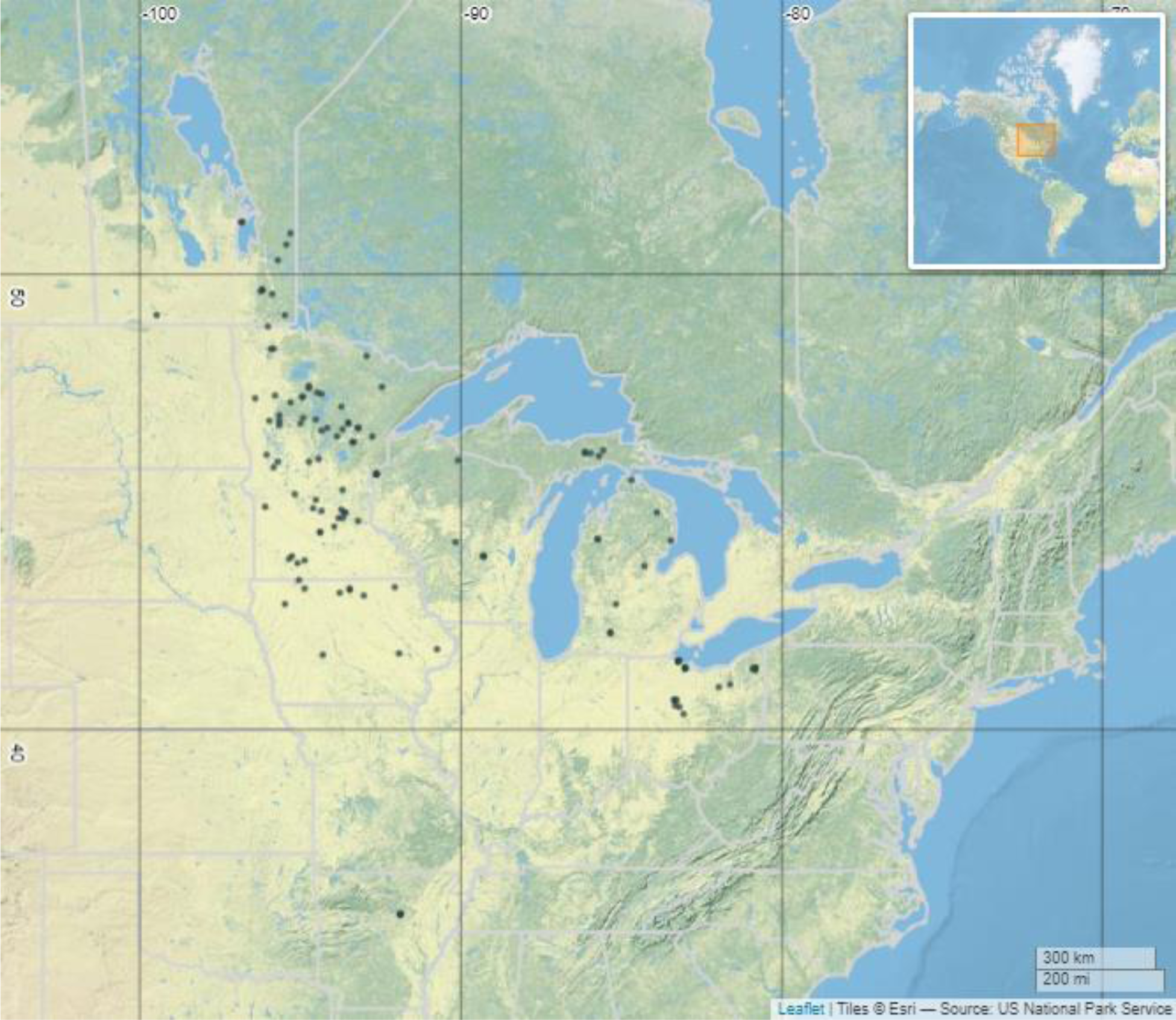
Capture locations of Interior Population trumpeter swans collared with GPS-GSM transmitters from July 2019–January 2022.

### Capture and Handling

We captured all swans during the definitive prebasic molt period (except for 4 captured in Arkansas using snares during the non-breeding period) when adult swans replace their remiges and are therefore flightless, using a combination of jon boats, airboats, step deck transom boats, and square-stern canoes. We primarily used long-tail mud motors (Powell Performance Fab, Hutchinson, Minnesota, USA) to navigate shallow wetlands where swans were located, though we captured some swans using boats with surface-drive motors (Gator-Tail, Loreauville, Louisiana, USA) or non-motorized canoes and kayaks. We hand-captured most swans using shepherd’s crook poles or large landing nets [41,42] and captured swans in Michigan using a shoulder-fired net gun. We predominantly targeted adult swans, which have higher survival rates than juveniles, to maximize the longevity of telemetry data collection [43].

We marked swans with two types of collars: 55-g neck collars with GPS-GSM transmitters incorporated into the collar housing (Model OrniTrack-N62 3G, Ornitela, Vilnius, Lithuania) and 140-g GPS transmitters (Model CTT-ES400, Cellular Tracking Technologies, [CTT], Rio Grande, New Jersey, USA) that were adhered to 64-mm neck collars (Haggie Engraving, Crumpton, Maryland, USA). All collar and transmitter types weighed <3% of the marked individual’s body weight, as per animal care requirements. Both types of neck collars contained a unique alpha-numeric code for visual identification. Swans captured in Michigan were fit with CTT collars and all other swans in the study were fit with Ornitela collars. All transmitters were programmed to collect GPS locations at 15-min intervals throughout the 24-hr daily period. In the United States, we leg-banded each swan with a U.S. Geological Survey butt-end aluminum band, and in Canada we banded each swan with a stainless steel locking-tab leg band. We determined sex via cloacal examination and assigned a breeding status to each swan at the time of capture depending on if a mate and one or more cygnets were present (‘breeder’ status), if a mate was present but no cygnets were observed (‘paired’ status), or if neither a mate nor cygnets were present (‘non-breeder’ status). Due to logistical constraints, we were unable to continuously monitor the breeding status of swans in subsequent years, and we retained the initial breeding status designation for all analyses. We are not aware of published information on the probability of trumpeter swans nesting in subsequent years. Previous research on mute swans (*Cygnus olor*) documented the probability of nesting in subsequent years at >90% [44,45]. We use the term breeding (e.g., breeding latitude, breeding territory) to indicate either breeding territory for successful breeders, or capture location for non-breeding swans (except for the 4 swans captured in Arkansas).

### Migration Phenology Classification

One of our key objectives was to accurately quantify IP trumpeter swan migration phenology. Given the size of the dataset (∼6M locations over 252 swan-years), we sought an efficient workflow to segment location data into periods of the annual cycle and to estimate migration metrics. Many migration segmentation approaches are based on spatiotemporal criteria that require subjective species-specific decisions, reducing reproducibility and limiting the potential to generalize to other studies, especially given the complexity of migratory behaviors that many species exhibit [46,47].

To more objectively quantify migration phenology, we used a model-driven approach with displacement from the breeding site used to segment the annual cycle into stationary periods that correspond with breeding, stopover, and non-breeding areas. We first calculated annual time-series of Net-Squared Displacement (NSD) values for each swan (referred to as a ‘swan-year’ dataset), using 1 July (a time of year when swans are relatively stationary) as a cutoff date between years for individuals with multiple years of GPS data and as an origin point for NSD values and then thinned datasets to contain a single averaged displacement value per day [16,48]. After excluding 11 swan-year datasets with <30 days of data, we iteratively fit a series of 7 intercept-only piecewise regression models to each time-series [48,49]. The syntax of each model corresponded to an increasing number of intercepts included (from 1–7) for average displacement values throughout the time series separated by breakpoints in time when the intercept values transitioned; therefore intercepts represent stationary segments in time corresponding to periods of the annual cycle, and breakpoints are the transitions between these segments that provide information on the timing of migration events such as autumn departure and spring arrival (Fig. 2).

**Figure 2:**
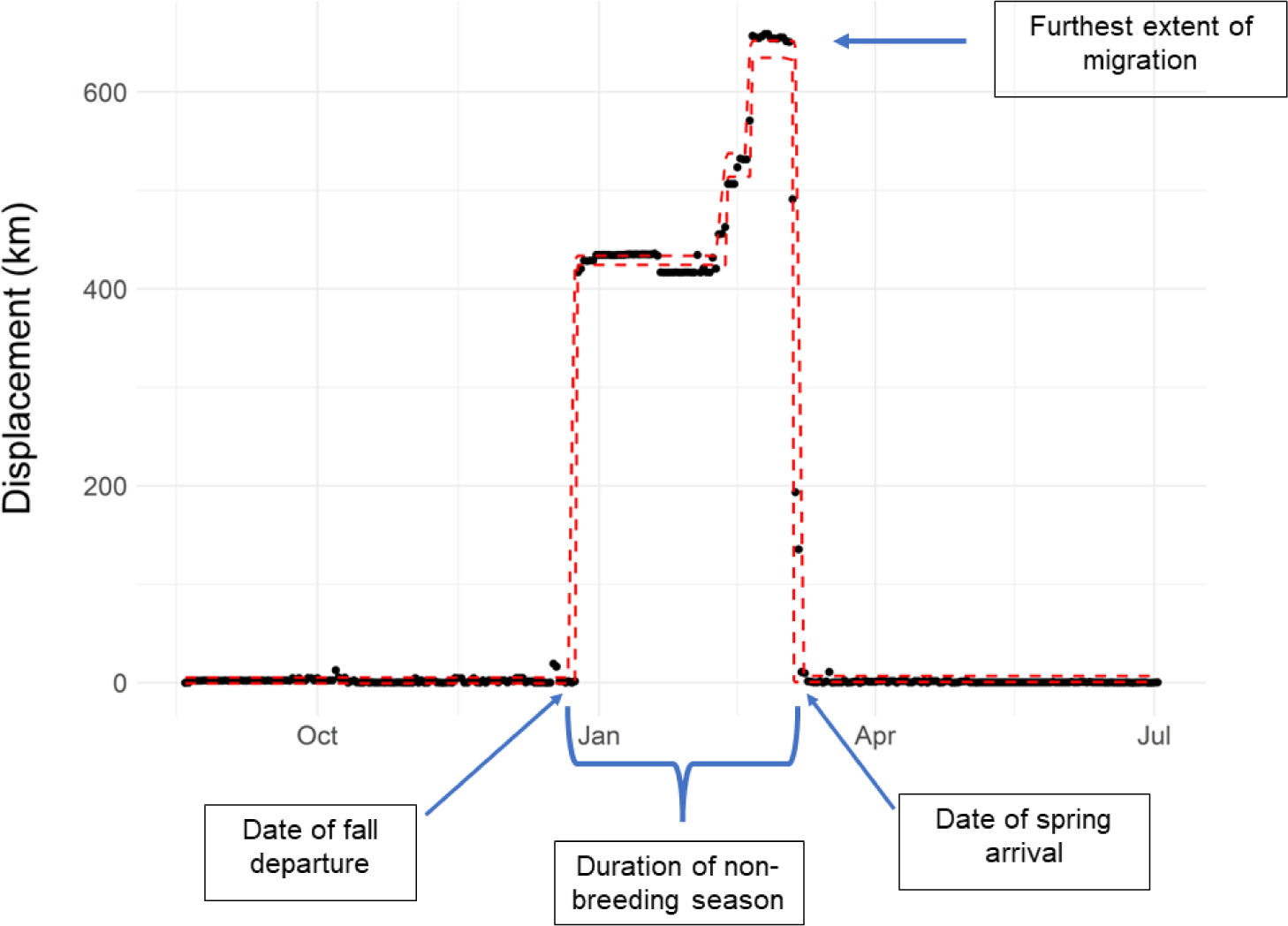
An example of extracting migration phenology metrics from a fitted piecewise regression model of an annual time-series of displacement values for an individual trumpeter swan.

We fit all piecewise regression models in JAGS [50] using the *mcp* package [49] in Program R version 4.0.2 [51], using its default priors and 15,000 iterations with a burn-in period of 10,000. We used the *future* R package to run all scripts in parallel via the Minnesota Supercomputing Institute using a partition with 24 cores and 50GB RAM per core [52]. We evaluated Markov-chain Monte Carlo (MCMC) convergence via the Gelman-Rubin statistic (*R_̂_*) and excluded models containing any parameters with a value of *R_̂_* >1.1 from further consideration [53]. Through extensive testing, we found that models with MCMC samples that failed to converge in distribution after 15,000 samples typically provided a poor fit to the data (e.g., single-intercept models despite multiple migration periods or models with multiple breakpoints during a stationary period). If all parameters in a model passed the *R_̂_* threshold, we evaluated model fit and predictive performance using leave-one-out cross-validation (LOO-CV) with Pareto smoothed importance sampling to estimate the Expected Log Predictive Density (ELPD), using the *loo* package [54,55]. The ELPD values reflect the ability of the model to predict the posterior density of withheld data. We used LOO-CV to choose the optimal number of breakpoints (and thereby segments that correspond to migratory periods) for each swan-year dataset.

We qualitatively inspected the visual fit of each best-supported model (based on ELPD) for each swan-year dataset and removed 11 (out of 241 total datasets) that had obvious poor fits such that information from the breakpoints and intercepts would not adequately describe annual migration phenology. To avoid confusing short-distance and temporary relocation, we also excluded all segments <2 km from the previous segment and all changepoints <2 days from the previous changepoint. We then extracted parameter values to represent the movement metrics of interest (Fig. 2). To focus on swans that made obvious movement away from breeding areas, we extracted autumn departure dates for individuals that had moved >100 km from breeding locations by 30 December (Fig. S2). We extracted spring arrival dates for individuals that moved >100 km from breeding locations during the non-breeding period and that returned within 30 km of their previous breeding territory (Fig. S2). We designated movement categories based on each swan’s annual migration extent (farthest distance moved from breeding/capture territory during the non-breeding period), and assigned categories of local movement (<25 km), regional migration (25–100 km), and long-distance migration (>100 km) for each swan-year dataset (Fig. S3).

### Migration Extent

An exploratory analysis of annual movement data suggested a strong linear association between breeding latitude and migration extent for one group of individuals, whereas others moved a much lesser extent, especially those at lower latitudes. It is possible that these patterns reflect two different migratory strategies with one segment of the population consistently migrating to lower latitudes and a second segment of the population using areas with open water closer to the location of their breeding site during the non-breeding period.

To describe these potential relationships between breeding latitude and migration extent, we used a 2-component mixture model (i.e., a model with two different groups of individuals, each following a different response pattern). Let *Y*_*ij*_ and *x*_*ij*_ represent the migration extent and breeding latitude, respectively, for individual *i* in year *j*, and let *z*_*i*_ be a latent variable representing the group membership of the *i*th individual. We assumed migration extent was linearly related to breeding latitude for individuals with *z*_*i*_ = 1 and was much lower and non-linearly related to breeding latitude for individuals with *z*_*i*_ = 0:

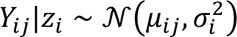

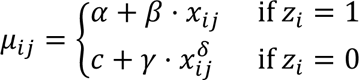

In addition, we assumed the variance of *Y*_*ij*_ depended on the latent state, *z*_*i*_.

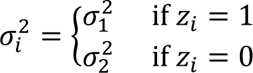

The latent state *z*_*i*_ (0 or 1) for assignment to a group (i.e., migration strategy) was modeled as:

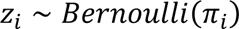

We used normal priors for *α* ∼ 𝒩(400,0.001), *β* ∼ 𝒩(150,0.001), and *γ* ∼ 𝒩(30,0.001), parameterized in terms of their mean and precision (1/*variance*), as is customary in JAGS-style notation. We used uniform priors for *c* ∼ *Uniform*(0,100), *δ* ∼ *Uniform*(0,30), *π* ∼ *Uniform*(0,1), *σ*_1_ ∼ *Uniform*(1,200), and *σ*_2_ ∼ *Uniform*(1,200). We used priors partially informed by the data for the intercepts of each group, *α* and *c*, based on the assumption that birds in group 1 migrate a greater extent than those in group 2.

We implemented the model in JAGS via the *R2jags* package using 3 MCMC chains with 30,000 iterations, a burn-in of 10,000, and a thinning rate of 10 [50,56]. We determined convergence of chains by assessing trace plots and using the Gelman-Rubin statistic with *R_̂_* <1.1[53].

### Migration Phenology

We visualized observations of migration phenology grouped by breeding status with boxplots showing the medians, inter-quartile ranges, and outliers. To test associations between predictor variables and migration phenology, we fit separate linear mixed models (LMM) using the *lme4* package to each response (autumn departure date, spring arrival date, and duration of non-breeding period), with a random intercept for each individual to account for repeated measures (i.e., multiple migrations by the same individual), and fixed effects for sex, breeding status, and breeding latitude [57]. We assessed model assumptions using the *performance* R package and plotted estimates of coefficients and associated 95% confidence intervals using the *sjPlot* R package [58,59]. To visualize the relationship between breeding latitude and each migration metric, we plotted estimated marginal effects from the linear mixed models using the *ggeffects* package [60]. We used the *emmeans* package to conduct post-hoc pairwise comparisons involving the categorical levels of breeding status (i.e., breeder, paired, non-breeder), with Tukey’s Honest Significant Difference (HSD) to adjust *p*-values and confidence intervals for each comparison [61]. We approximated the degrees of freedom for the HSD test of the spring arrival LMM using the Kenward–Roger method [62].

## Results

We deployed 113 collars with GPS-GSM transmitters on 126 trumpeter swans (including 13 redeployments using collars recovered from mortalities) from July 2019 to August 2022, resulting in 252 unique ‘swan-year’ telemetry datasets (Fig. S1). Of the 126 swans, 78 were female and 48 were male; 73 were breeding adults (cygnets present), 22 were adults with mates present but no cygnets observed at time of capture, 24 were non-breeding adults captured while in large groups, and six were cygnets at the time of capture (however, data from cygnets were not used in analyses).

### Migration Extent

Annual movements of IP trumpeter swans were highly variable. After filtering out 31 swan-year datasets from 27 individuals with incomplete coverage to estimate annual movement trends, 59% (*n*=68) of remaining IP swans (*n*=116 individuals) made seasonal movements to distant non-breeding areas (long-distance migration defined as moving >100 km from breeding site during the non-breeding period), 16% (*n*=19) underwent regional migration (25-100 km), 19% (*n*=22) exhibited local movements (<25 km), and 6% (*n*=7) exhibited multiple migration strategies across different years: regional and long-distance (*n*=3), and local and regional (*n*=4). The mixture model applied to migration extent classified all individuals to a group with high probability, so it was unnecessary to set a user-defined threshold to determine group assignment. There was a strong association between breeding latitude (mostly between 43 and 53 degrees North latitude) and the extent of migration (Fig. 3). Yet, many swans with breeding sites between 40 and 45 degrees North latitude exhibited minimal movement during the non-breeding period. We considered these individuals to exhibit local seasonal movements, with most of these swans leaving their breeding territory or increasing their overall space use during the non-breeding period, likely to increase access to ice-free water or to food.

**Figure 3:**
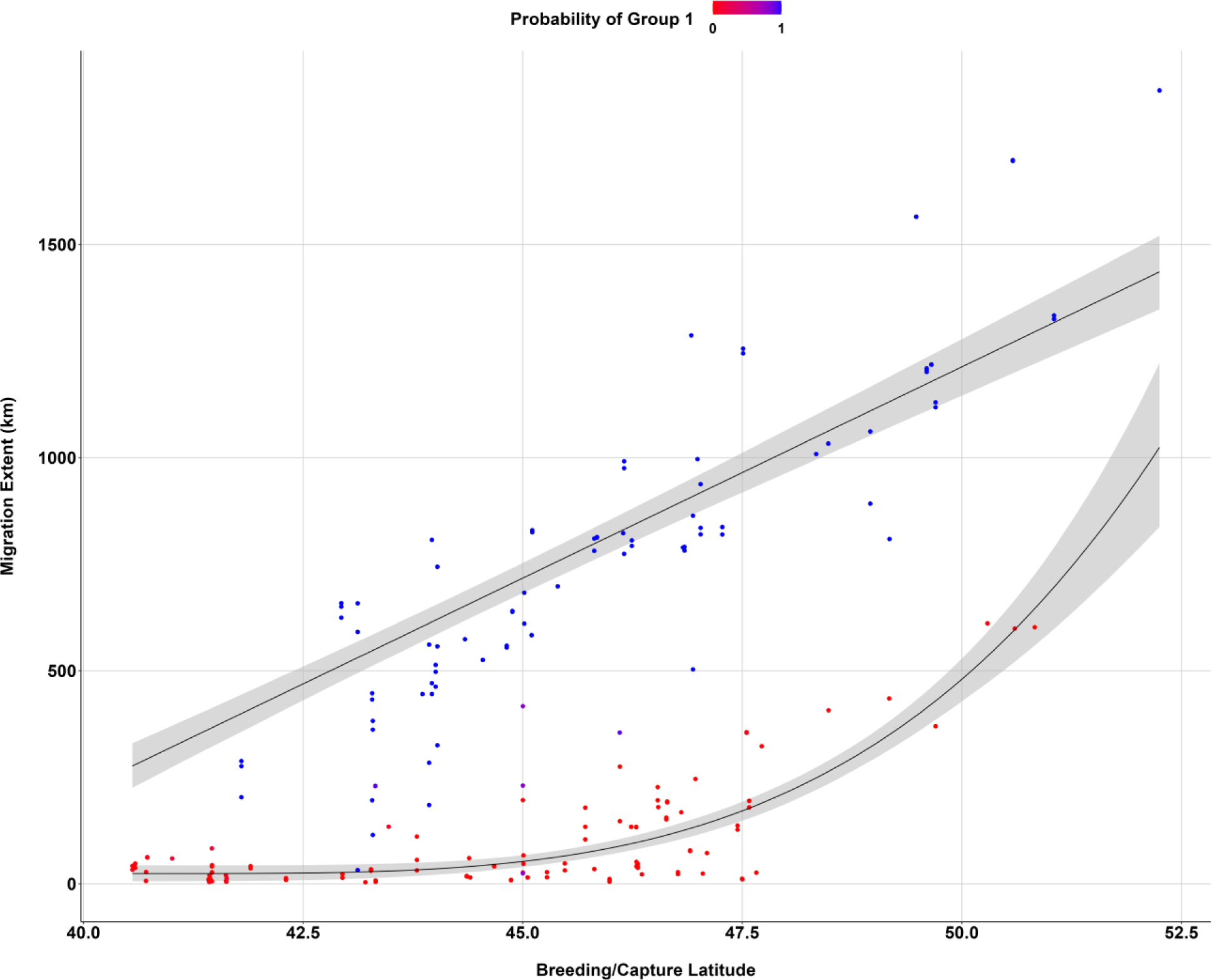
Migration extent versus breeding/capture latitude with color indicating the probability of assignment to one of two migration strategies within a 2-component mixture model describing the relationship between latitude and migration extent. Gray areas depict the 95% credible intervals for each strategy.

### Migration Phenology

#### Autumn Departure

The average autumn departure date from breeding territories across years for all long-distance (>100 km) migrants was 7 November with a standard deviation of 25 days, and yearly averages ranged from 31 October to 17 November (Table S1, Table S2). Long-distance migrants at higher latitudes left territories earlier in autumn than swans at lower latitudes. On average, swans left 4 days earlier (95% CI = −6.2– −1.9) for every increase of 1 degree of breeding latitude (corresponding to about 111 km; Fig. 4, Table S10). Estimated autumn departure dates of breeding swans averaged 18 days (95% CI = 2.9–33.9) later than non-breeders and 6 days (95% CI = −8.7–20.9) later than paired swans (Fig. 4, Fig. 6, Table S3, Table S11).

**Figure 4:**
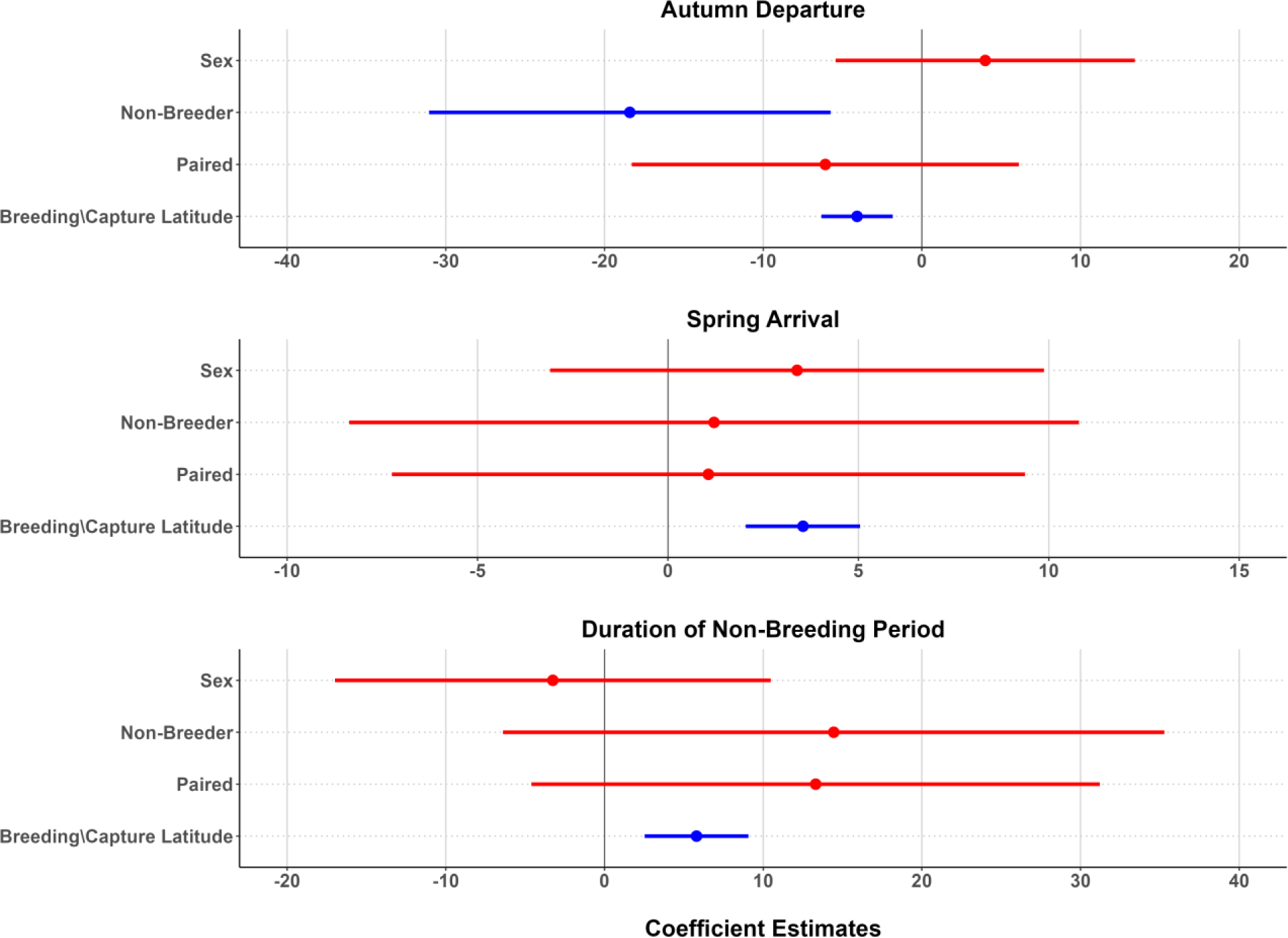
Estimates and 95% confidence intervals for model coefficients from linear mixed effects models fit to autumn departure, spring arrival, and duration of non-breeding period. The covariate for sex represents the difference between male and female (the reference category). The coefficients for non-breeder and paired contrast these categories with breeder (the reference category). Blue results represent statistically significant coefficient estimates and red results represent coefficient estimates that were not statistically significant.

**Figure 5:**
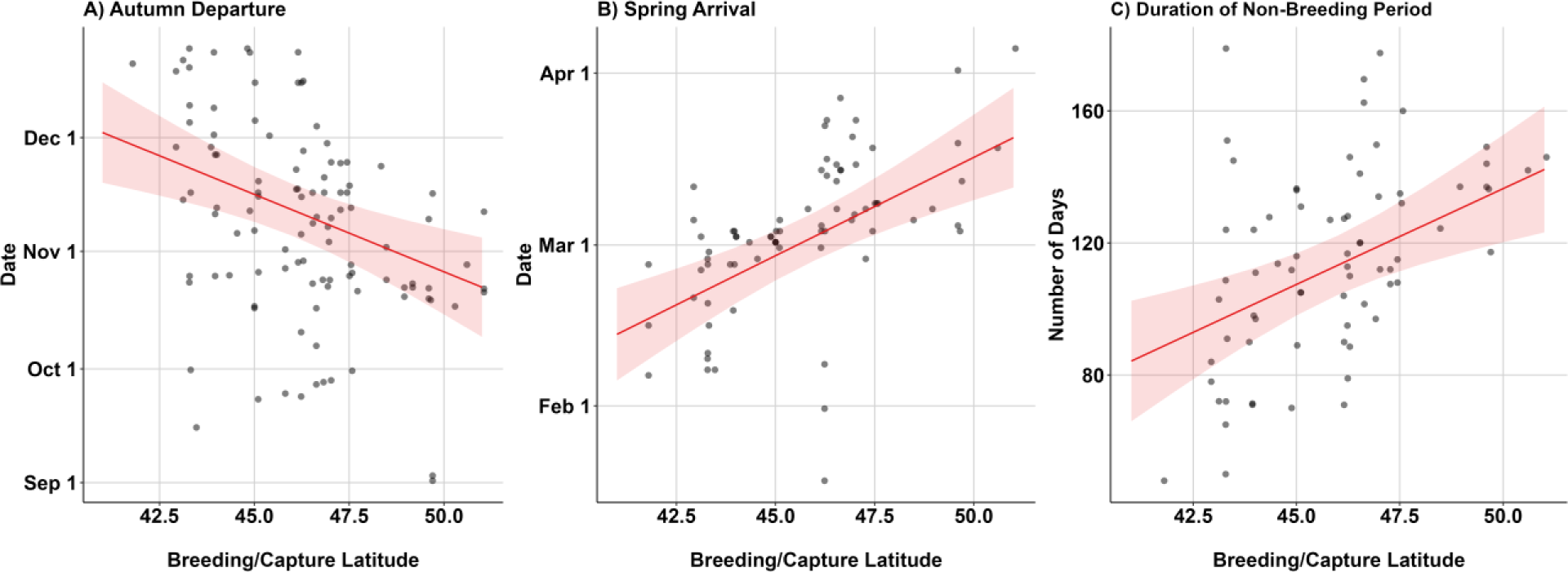
Marginal effect plots for the relationships between breeding/capture latitude and autumn departure (A), spring arrival (B), and duration of non-breeding period (C). Red lines and shaded areas depict the predicted values and 95% confidence intervals, respectively. Points shown are the observed migration phenology metrics.

**Figure 6:**
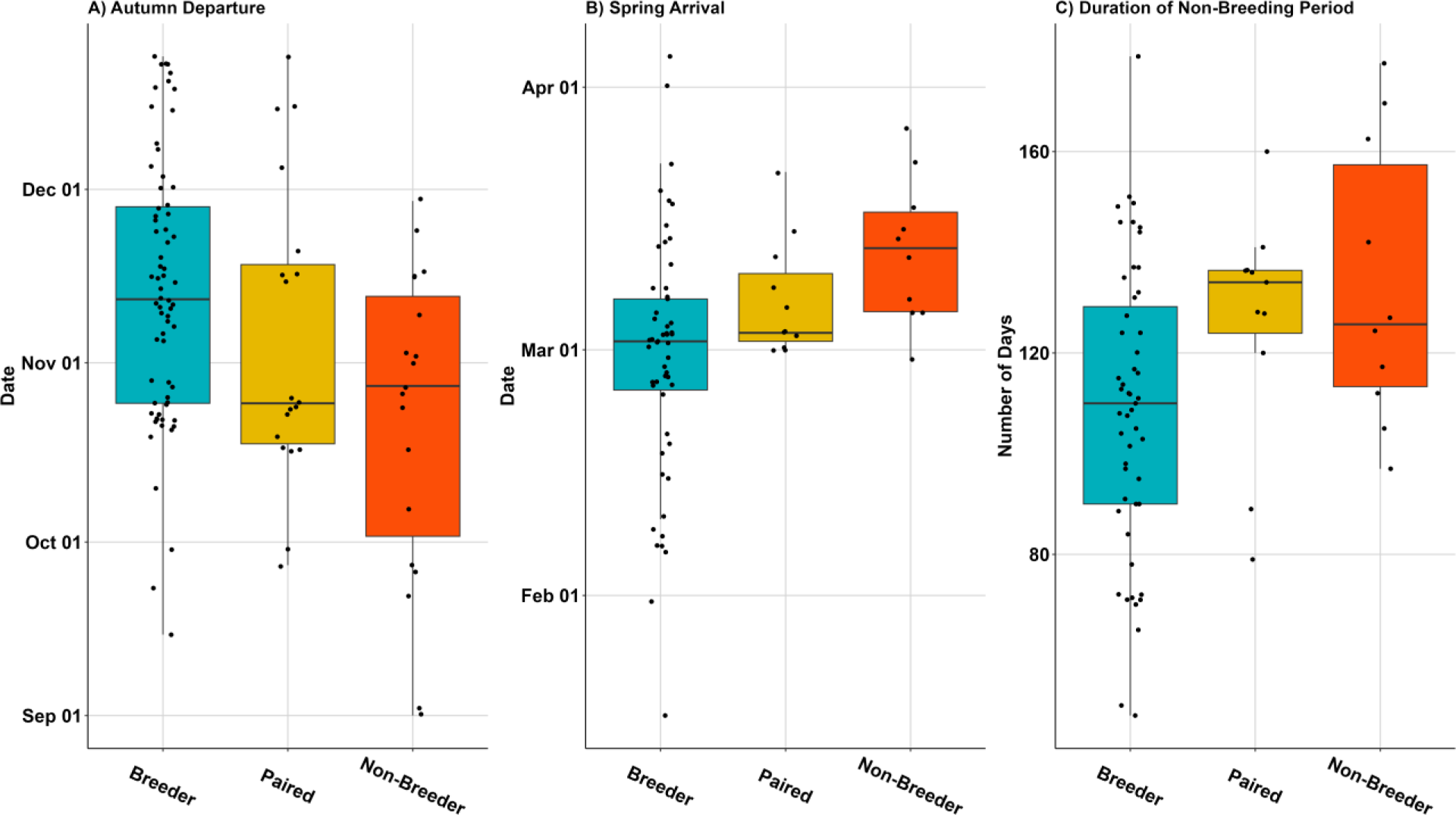
Migration phenology dates by breeding status (breeder, paired, and non-breeder). Boxes bound the 25th and 75th percentiles, solid lines within the boxes indicate the median, lines extend to 1.5 times the interquartile range, and points correspond to individual swan-years.

#### Spring Arrival

The average spring arrival date across years for all long-distance (>100 km) migrants was 4 March with a standard deviation of 15 days, and yearly averages ranged from 2 March to 6 March (Table S4, Table S5). Swans that returned to within 30 km of their previous territory arrived earliest at southern latitudes. On average, swans arrived 5 days (95% CI = 3.9–6.4) later for every increase of 1 degree of breeding latitude (Fig. 4, Table S10). Differences in spring arrival among breeding categories were minimal (Fig. 6, Table S6), and confidence intervals included zero for all two-way comparisons (breeding–non-breeder, *β* = −1.2, 95% CI = −12.9–10.5; breeder–paired, *β* = −1.1, 95% CI = −11.2–9.1; non-breeder–paired, *β* = −0.1, 95% CI = −12.9–13.2; Fig. 4, Table S12).

#### Duration of Non-Breeding Period

The average duration of the non-breeding period for swans that had estimated autumn departure dates and also spring arrival dates the following year was 115 days with a standard deviation of 29 days, and yearly averages ranged from 99 days to 119 days (Table S7, Table S8). On average, long-distance migrants at higher latitudes had longer non-breeding periods, with every increase of 1 degree of breeding latitude equating to 6 additional days away from the breeding territory (95% CI = 2.6–8.9; Fig. 4, Table S10). Long-distance migrants classified as breeders tended to have shorter non-breeding periods (Fig. 6, Table S9), but confidence intervals included zero for all two-way comparisons (breeder–non-breeder, *β* = −14.5, 95% CI = −40.0–11.1; breeder–paired, *β* = −13.3, 95% CI = −35.3–8.7; non-breeder–paired, *β* = −1.1, 95% CI = −27.3–29.6; Fig. 4, Table S13)

## Discussion

Annual movements of IP swans were highly variable in their extent and timing, with a continuum of movements exhibited each year. Much of this variability was related to factors tied to natural history demands (e.g., breeding status) and response to environmental conditions (e.g., through associations with breeding latitude). It is not immediately clear why IP swans seemed to exhibit multiple migration strategies even at similar latitudes where the effects of environmental conditions (e.g., temperature, precipitation) should also be similar, but other factors such as social dynamics, genetic lineage, and site-specific differences in the availability of open water and food likely play an important role. Fine-scale spatial variation, especially related to open water, may have a large influence on determining which swans undergo long-distance migration and which can spend the non-breeding period at higher latitudes.

The migration phenology of IP trumpeter swans appears similar to that of other long-lived migratory birds, with breeders leaving later in the autumn than non-breeders and spending less time away from the breeding territory during the non-breeding period [63–65]. Autumn departure, spring arrival, and duration of non-breeding period were also highly correlated with breeding latitude (Fig. 4), likely driven by variable access to open water. Breeding latitude was also associated with migration extent during the non-breeding period (Fig. 3). Individuals that bred in the northern part of the IP range made relatively long-distance autumn migrations. Given the severity of winter conditions at higher latitudes, options for accessing open water created by currents on rivers or anthropogenic influences (e.g., below dams, on lakes with aerators) were likely not near to breeding locations. Autumn migration distances of swans breeding at mid-latitudes were variable, with some swans moving relatively long distances, whereas others exhibited only local or regional movements. These latter swans likely remained near their breeding locations during the non-breeding period if they were able to find open water and had access to food. Swans breeding at lower latitudes remained close to their breeding locations year-round, as the local environmental conditions likely continued to provide open water and access to food.

Several other factors may influence movements and migratory patterns, including transmission of migratory information between generations genetically and through social learning [66]. For some species with short lifespans or limited parental care (e.g., songbirds, raptors), migration is considered innate and primarily genetically controlled, based on observations of individuals that complete their first migrations independently without parents or other conspecifics to guide them [67]. However, for species with long generation times and extended periods of parental care, such as trumpeter swans, social learning typically also plays an important role in forming migration strategies [9]. Collective knowledge has been shown to accumulate over generations to drive migration patterns and improve efficiency in flocking species with socially learned migration behaviors [12]. In reintroduced populations of whooping cranes (*Grus americana*) that were initially trained (i.e., learned) to migrate by following aircraft, migratory efficiency of flocks rapidly increased when older individuals were present [9,68]. Thus, experience and familiarity with the landscape also likely play important roles in determining migratory movements. For swans, in particular, knowledge of sites with access to open water and food may be important for allowing swans to ‘escape’ migration in more northern latitudes, and this knowledge may be passed down through generations.

IP swans are unique because their reintroduction to central North America occurred relatively recently (<50 years) from a mix of source populations with various migratory strategies [69]. Thus, there may exist heterogeneity within the population in terms of genetic makeup that influences swan movements. Although previous studies have considered genetic differences of IP source populations using microsatellites, contemporary research methods (i.e., population genomics) are needed to better evaluate the genetic structure of the IP as it relates to other trumpeter swan populations [19,69]. Continued monitoring is also necessary to determine whether population-level migration characteristics (e.g., annual proportion of IP swans that migrate) are still in flux or if conditions have stabilized akin to the environmental threshold model, in which certain parts of the population are obligate residents or migrants and annual environmental conditions determine the migration threshold that dictates the outcome of facultative migrants, a population-level paradigm thought to be maintained predominantly through genetic variation [70]. Ultimately, our study provides a snapshot in time of current IP trumpeter swan migration patterns ∼50 years following initial reintroduction efforts, and population-level migration patterns may still be stabilizing and exhibiting variation based on annual environmental conditions.

In addition, the history of IP reintroduction includes many years of anthropogenic influences, from both wildlife managers and private citizens, that have likely affected migration patterns, and it is not well understood to what extent these impacts may be passed down through generations [15]. Some of these influences include intentional feeding during the non-breeding period at sites of high IP swan density and lake aerators that keep lakes ice-free throughout the winter to prevent winter kill of fish [71–73]. Although supplemental feeding has been discontinued by state and provincial management agencies throughout the IP distribution, the practice continues with some private citizens, and potential effects include not only curtailment of migration but also increased risk of pathogen transmission [74]. There is also increasing evidence that many local groups of swans have discovered field feeding as an additional strategy to acquire resources, and this knowledge likely affects migratory behavior, though additional research is needed to quantify the spatial extent, intensity, and timing of field-feeding in trumpeter swans. [75,76].

## Conclusions

Knowledge of current IP swan migration patterns can help inform IP swan conservation by providing current information on the limits of the non-breeding period range and quantifying variability in migration strategies. Under current conditions, non-breeding-period habitat for swans occurs in all but the most northerly portions of the IP breeding distribution. Managers may conserve habitat during the non-breeding period for both long-distance migrants and residents that remain on their breeding territories year-round while also anticipating the effects of future climate scenarios [77]. It is not clear how changing climate conditions will influence future IP swan migration strategies, but currently, multiple migration strategies exist within the IP, providing a range of behaviors as the basis for adaptation to changing conditions that may help offset potential negative effects of asynchrony between migration phenology and environmental conditions [78]. The variability present in IP annual movements may position IP swans to quickly adapt to conditions resulting from changing climate.

## Availability of data and materials

All R code used in this analysis are available at https://github.com/dwwolfson/swan_migration_manuscript. Data used to support this study are available from the corresponding author upon request.

## Declarations

### Ethics approval and consent to participate

Protocols for capturing and marking trumpeter swans in the United States were approved by the University of Minnesota Animal Care and Use Committee (protocol no. 1905-37072A), the U.S. Fish and Wildlife Service (Research & Monitoring Special Use Permits no. K-10-001, M-20-002, and M-21-014), the U.S. Geological Survey Bird Banding Laboratory (Federal Bird Banding Permit no. 21631) and state-specific permits approved by each state wildlife agency involved. All captures and marking of trumpeter swans in Manitoba were conducted under Federal Scientific Permit to Capture and Band Migratory Birds (no. 10271), Federal Animal Care Committee approval (project 20FB02), Provincial Species at Risk Permit (no. SAR20012), and Provincial Park Permit (no. PP-PHQ-20-016).

### Consent for publication

Not Applicable

### Authors’ contributions

DWW, JRF, and DEA designed the methodology; DWW, RTK, ABT, LK, BK, TF, TH, SM, TM, DF, and DEA collected the data; DWW analyzed the data; DWW wrote the first draft of the manuscript, which was revised primarily by DWW, JRF, and DEA. All authors reviewed the manuscript and gave final approval for publication.

## Acknowledgments

We thank Dustin Arsnoe, Rob Batterson, Laurie Brown, Tom Cooper, Steve Cordts, Bruce Davis, Victoria Drake, Anthony Duffiney, Jennifer Fredrickson, Matthew Garrick, David Hoffman, Steven Hogg, Joel Huener, John Hummel, Gary Ivey, Doug McArthur, Ciara McCarty, Luke Naylor, Karen Norris, Mike North, Karen Rowe, Rachel Ruden, Jess Schmidt, Brendan Shirkey, Nikki Smith, Erik Thorson, Geoff Westerfield, Sara Zaleski, and Ed Zlonis for assistance with fieldwork. We thank Barb Avers, Frank Baldwin, Kent Bekker, Wayne Brininger, Tom Cooper, Peter David, Walt Ford, David Luukkonen, Hattie Saloka for assistance with project logistics. We thank Rachel Vanausdall and Doug Johnson for constructive comments that improved the manuscript. Any use of trade, product, or firm names is for descriptive purposes only and does not imply endorsement by the U.S. Government.

