## Supplemental Figures and Tables for "High variability of migration strategies in a re-established Trumpeter Swan population"

[8] Three Rivers Park District

[9] Cleveland Metroparks Zoo

[10] Trumpeter Swan Society

[11] Minnesota Department of Natural Resources

[12] U.S. Geological Survey, Minnesota Cooperative Fish and Wildlife Research Unit

[13] University of Minnesota

#### Supplemental Figures

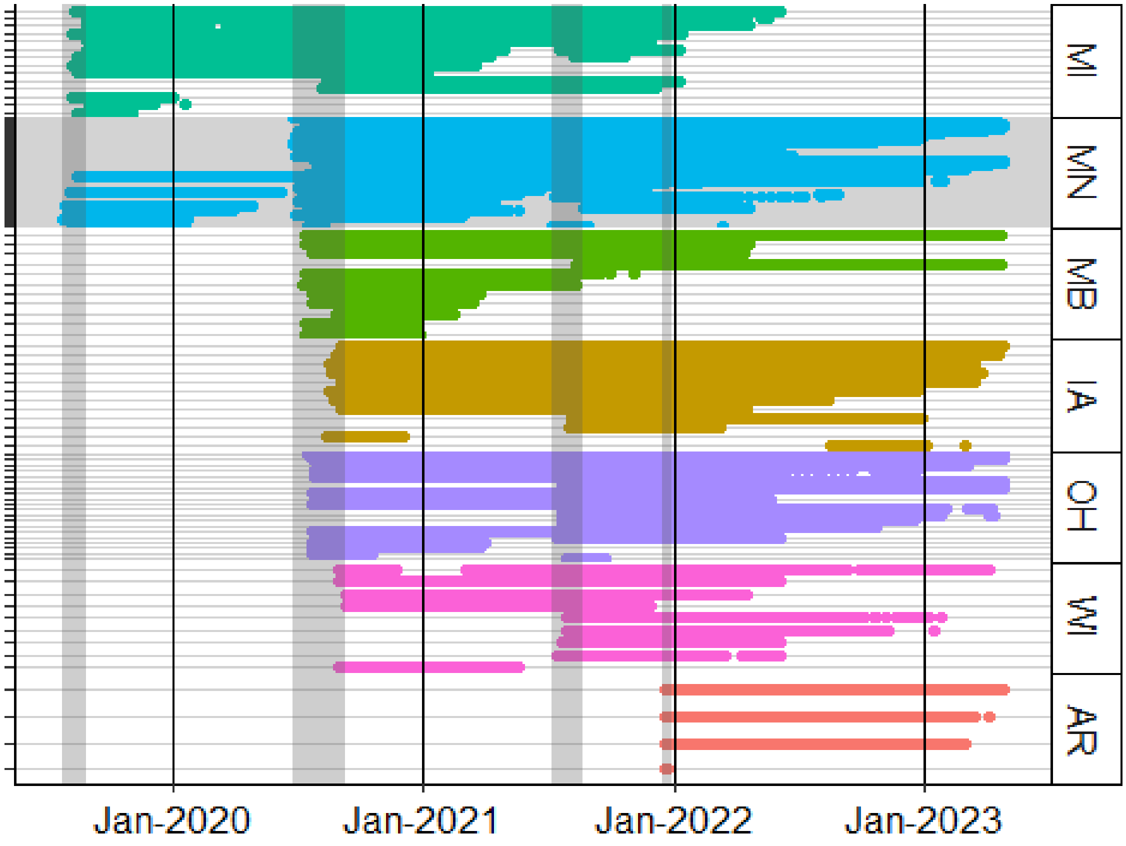


Figure S1. An overview of GPS telemetry data received from all collared IP trumpeter swans. Each line represents the period of data collection from a single collared swan. The grey regions indicate periods of collar deployment. The black lines are 1 January of each year. Number of deployments (including redeployments) by state/province are: Michigan (MI, *n*=14), Minnesota (MN, *n*=56), Manitoba (MB, *n*=11), Iowa (IA, *n*=12), Ohio (OH, *n*=20), Wisconsin (WI, *n*=9), and Arkansas (AR, *n*=4).

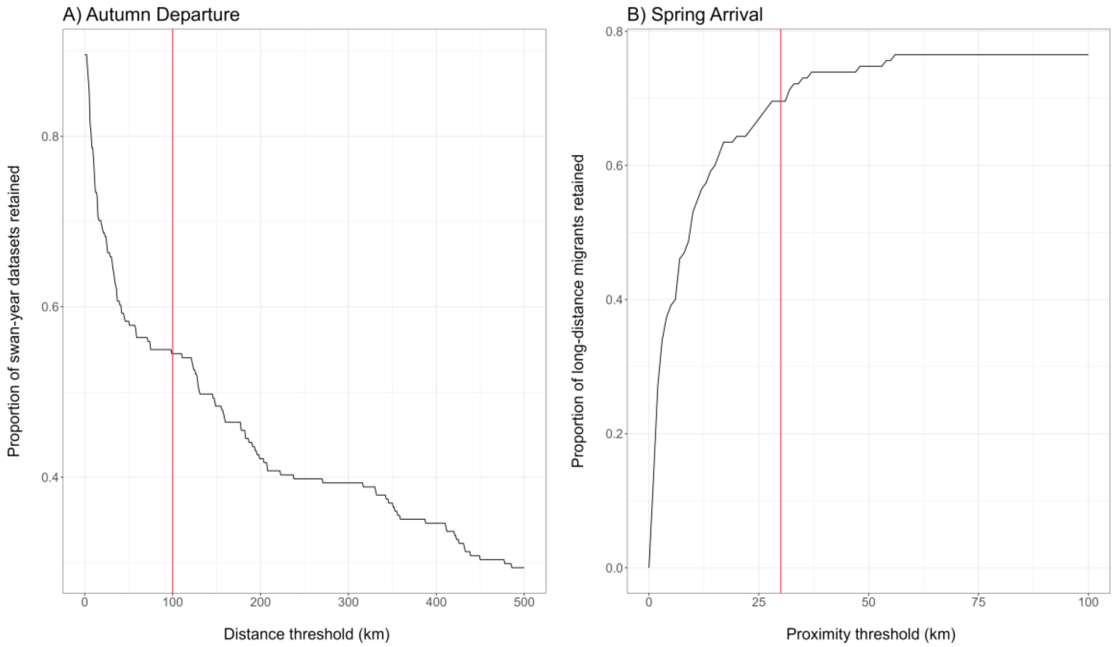


Figure S2. Sensitivity analyses for the thresholds used as cutoffs for whether to estimate migration timing (autumn departure and spring arrival). (A) The proportion of swan-year datasets that satisfy distance threshold cutoffs (0–500 km) as well as other rules (e.g., ignoring changepoints <2 days and segments <2 km from each other, potential departure before 30 Dec) for being considered a 'long-distance' migrant, and estimating an autumn departure date. The red line indicates the threshold of 100 km used in the analysis. (B) The proportion of swan-year datasets that satisfy a proximity threshold cutoff (0-100 km), defined as the required proximity to the previous year's breeding/capture territory (i.e., the distance from the previous year territory that the swan must be within the following year), as well as other rules (e.g., previously satisfied criteria for estimating an autumn departure, potential arrival after 30 Dec) used to estimate a spring arrival date. The red line indicates the threshold of 30 km used in the analysis.

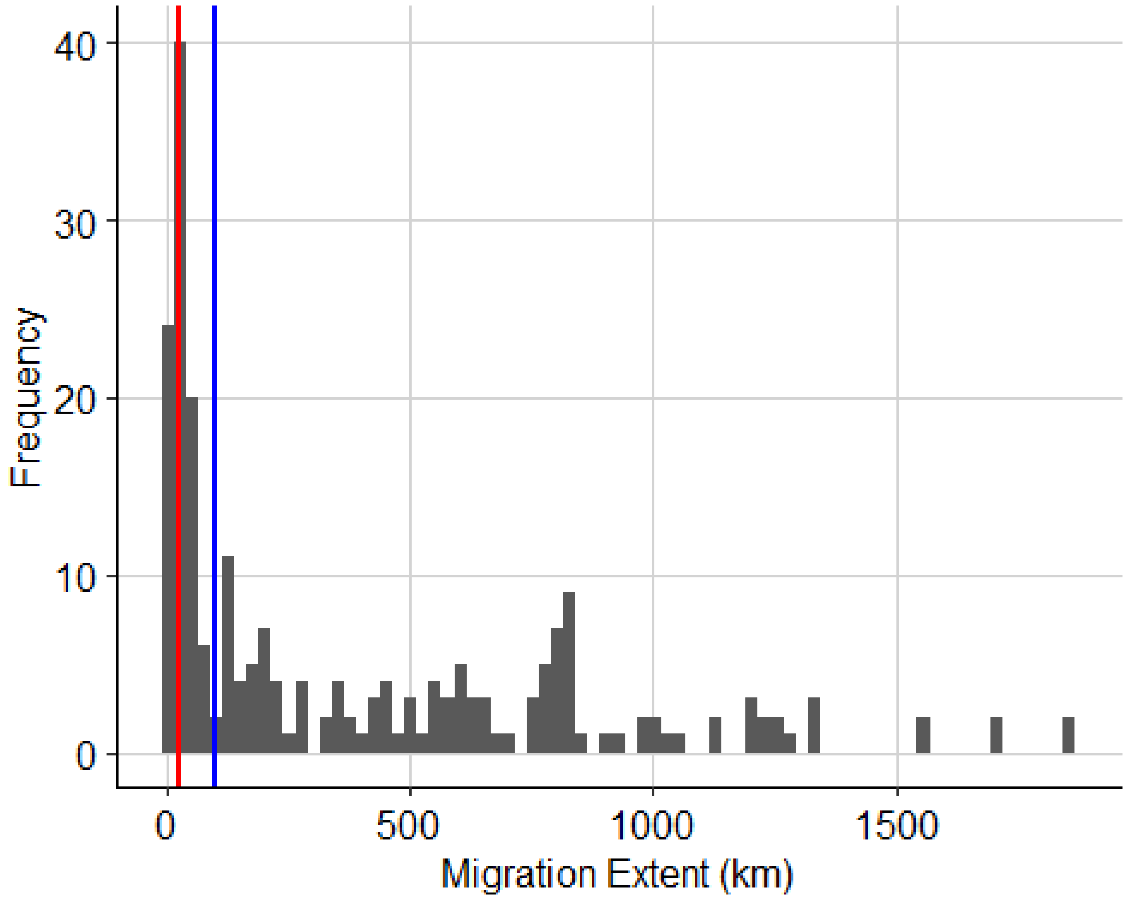


Figure S3. Histogram of the migration extent (farthest distance from breeding territory during the non-breeding period, i.e., maximum annual displacement segment from piecewise regression model) for all 221 swan-year datasets. The distance thresholds used to define categories of migration (local movements = 0-25 km, regional migration = 25-100 km, long-distance migration >100 km) are shown with the red (25 km) and blue (100 km) lines.

### Supplemental Tables

#### Migration Phenology Summary Statistics

##### Autumn departure

We estimated autumn departure dates for all swans that traveled >100km from the breeding/capture territory by 30 December.

Table S1. Compiled migration phenology of all autumn departures from 2019–2022.

| Total Swans Tracked | Number of Long-Distance Migrants | Number of Fall Departure Events | Average Autumn Departure | Standard Deviation (days) | Earliest Departure | Latest Departure |
|---|---|---|---|---|---|---|
| 122 | 71 | 117 | November 07 | 25 | September 01 | December 24 |

Table S2. Yearly summaries of migration phenology of autumn departures from 2019–2022.

| Year | Total Swans Tracked | Number of Long-Distance Migrants | Average Autumn Departure | Standard Deviation (days) | Earliest Departure | Latest Departure |
|---|---|---|---|---|---|---|
| 2019 | 17 | 7 | October 31 | 7 | October 21 | November 08 |
| 2020 | 82 | 49 | November 02 | 28 | September 01 | December 24 |
| 2021 | 86 | 38 | November 09 | 24 | September 15 | December 23 |
| 2022 | 44 | 23 | November 17 | 18 | October 07 | December 20 |

Table S3. Autumn departure dates of long-distance migrants by breeding status.

| Breeding Status | Total Swans Tracked | Number of Long-Distance Migrants | Average Autumn Departure | Standard Deviation (days) | Earliest Departure | Latest Departure |
|---|---|---|---|---|---|---|
| Breeder | 70 | 36 | November 13 | 23 | September 15 | December 24 |
| Non-Breeder | 22 | 12 | October 22 | 27 | September 01 | November 29 |
| Paired | 21 | 14 | November 04 | 26 | September 27 | December 24 |

**Spring arrival**

We estimated spring arrival for all swans that traveled >100 km from the breeding/capture territory during the non-breeding period, left their territory by 30 December, and returned to <30 km of their previous year's territory.

Table S4. Compiled migration phenology of all spring arrivals from 2020–2023.

| Total Swans Tracked | Number of Long-Distance Migrants | Number of Spring Arrival Events | Average Spring Arrival | Standard Deviation (days) | Earliest Arrival | Latest Arrival |
|---|---|---|---|---|---|---|
| 122 | 51 | 84 | March 04 | 15 | January 18 | April 11 |

Table S5. Yearly summaries of migration phenology of spring arrivals from 2020–2023.

| Year | Total Swans Tracked | Number of Long-Distance Migrants | Average Spring Arrival | Standard Deviation (days) | Earliest Arrival | Latest Arrival |
|---|---|---|---|---|---|---|
| 2020 | 17 | 4 | March 02 | 18 | February 08 | March 23 |
| 2021 | 82 | 35 | March 03 | 8 | January 31 | March 23 |
| 2022 | 86 | 33 | March 06 | 19 | January 18 | April 11 |
| 2023 | 44 | 12 | March 02 | 20 | February 06 | April 08 |

Table S6. Spring arrivals of long-distance migrants by breeding status.

| Breeding Status | Total Swans Tracked | Number of Long-Distance Migrants | Number of Spring Arrival Events | Average Spring Arrival | Standard Deviation (days) | Earliest Arrival | Latest Arrival |
|---|---|---|---|---|---|---|---|
| Breeder | 70 | 29 | 54 | February 29 | 15 | January 18 | April 05 |
| Non-Breeder | 22 | 8 | 10 | March 12 | 9 | February 29 | March 27 |
| Paired | 21 | 8 | 11 | March 06 | 7 | March 01 | March 22 |

**Duration of non-breeding period**

We estimated duration of non-breeding period for all swans that had an autumn departure (i.e., traveled >100 km from territory) followed by a spring arrival the following year. This migration metric represents the span of time absent from the breeding/capture territory during the non-breeding period, and is calculated by the difference in days between spring arrival and the previous year's autumn departure.

Table S7. Compiled duration of non-breeding period for all swans from 2019–2023.

| Total Swans Tracked | Number of Long-Distance Migrants | Number of Annual Cycles | Average Duration of Non-breeding Period (days) | Standard Deviation (days) |
|---|---|---|---|---|
| 122 | 49 | 78 | 115 | 29 |

Table S8. Yearly summaries of duration of non-breeding period from 2019–2020 until 2022–2023.

| Year | Total Swans Tracked | Number of Long-Distance Migrants | Average Duration of Non-breeding Period (days) | Standard Deviation (days) |
|---|---|---|---|---|
| 2020 | 17 | 4 | 119 | 21 |
| 2021 | 82 | 34 | 118 | 29 |
| 2022 | 86 | 28 | 117 | 28 |
| 2023 | 44 | 12 | 99 | 33 |

Table S9. Summaries by breeding status of duration of non-breeding period from 2019–2020 until 2022–2023.

| Breeding Status | Total Swans Tracked | Number of Long-Distance Migrants | Average Duration of Non-breeding Period (days) | Standard Deviation (days) |
|---|---|---|---|---|
| Breeder | 70 | 29 | 109 | 29 |
| Non-Breeder | 22 | 8 | 133 | 28 |
| Paired | 21 | 8 | 126 | 23 |

### Migration phenology model output

#### Linear mixed models

Table S10. Model summaries from 3 linear mixed models fit using the 3 migration metrics (autumn departure, spring arrival, and duration of non-breeding period) as the response. 95% confidence intervals for each coefficient are shown in brackets.

| | Autumn Departure | Spring Arrival | Duration of Non-breeding Period |
|---|---|---|---|
| (Intercept) | 323.701*** | 84.549* | -153.650* |
| | [219.816, 427.586] | [15.781, 153.317] | [-303.982, -3.317] |
| Sex | 3.993 | 3.389 | -3.260 |
| | [-5.440, 13.427] | [-3.097, 9.874] | [-16.994, 10.474] |
| Breeder – Non-Breeder | -18.404** | 1.211 | 14.446 |
| | [-31.058, -5.751] | [-8.372, 10.794] | [-6.395, 35.288] |
| Breeder – Paired | -6.083 | 1.063 | 13.308 |
| | [-18.281, 6.115] | [-7.250, 9.375] | [-4.602, 31.217] |
| Breeding/Capture Latitude | -4.082*** | 3.545*** | 5.802*** |
| | [-6.333, -1.830] | [2.045, 5.045] | [2.530, 9.074] |
| SD (Intercept swan_ID) | 7.240 | 7.432 | 6.137 |
| SD (Observations) | 21.615 | 7.981 | 24.752 |
| Num.Obs. | 107 | 75 | 72 |
| R2 Marg. | 0.228 | 0.338 | 0.265 |
| R2 Cond. | 0.306 | 0.645 | 0.308 |
| ICC | 0.1 | 0.5 | 0.1 |

+ p < 0.1, * p < 0.05, ** p < 0.01, *** p < 0.001

**Pairwise contrasts of migration timing by breeding status, adjusted using Tukey's HSD**

Table S11. Pairwise contrasts of autumn departure dates by long-distance migrants considered by breeding status.

| Contrast | Estimate | SE | Degrees of freedom | Lower 95% CL | Upper 95% CL | t ratio | p value |
|---|---|---|---|---|---|---|---|
| Breeder - Non-Breeder | 18.40 | 6.43 | 56.76 | 2.93 | 33.88 | 2.86 | 0.02 |
| Breeder - Paired | 6.08 | 6.18 | 65.44 | -8.74 | 20.90 | 0.98 | 0.59 |
| Non-Breeder - Paired | -12.32 | 7.72 | 65.81 | -30.83 | 6.19 | -1.60 | 0.25 |

Table S12. Pairwise contrasts of spring arrival dates by long-distance migrants considered by breeding status.

| Contrast | Estimate | SE | Degrees of freedom | Lower 95% CL | Upper 95% CL | t ratio | p value |
|---|---|---|---|---|---|---|---|
| Breeder - Non-Breeder | -1.21 | 4.82 | 45.60 | -12.88 | 10.45 | -0.25 | 0.97 |
| Breeder - Paired | -1.06 | 4.18 | 43.80 | -11.20 | 9.07 | -0.25 | 0.96 |
| Non-Breeder - Paired | 0.15 | 5.39 | 45.13 | -12.90 | 13.20 | 0.03 | 1.00 |

Table S13. Pairwise contrasts of duration of non-breeding period by long-distance migrants considered by breeding status.

| Contrast | Estimate | SE | Degrees of freedom | Lower 95% CL | Upper 95% CL | t ratio | p value |
|---|---|---|---|---|---|---|---|
| Breeder - Non-Breeder | -14.45 | 10.56 | 46.78 | -40.01 | 11.11 | -1.37 | 0.37 |
| Breeder - Paired | -13.31 | 9.06 | 44.53 | -35.28 | 8.67 | -1.47 | 0.32 |
| Non-Breeder - Paired | 1.14 | 11.76 | 48.12 | -27.29 | 29.57 | 0.10 | 0.99 |
